# Design and testing of a PCR assay to quantify coliform bacteria in drinking water

**DOI:** 10.1101/2024.09.06.611718

**Authors:** Claire Thom, Graeme Moore, Paul Weir, Cindy J Smith

## Abstract

Here we present an enhanced *lacZ* qPCR assay targeting β-galactosidase, encoded by lacZ, to assess coliforms in drinking water. Coliform bacteria are a diverse group of over 80 species, commonly found in enteric environments, and act as indicators of water supply contamination. Due to the significant variation among coliforms, and the size of the group designing a single target was challenging, as previous studies have demonstrated. To address this, a coliform *lacZ* sequence database was created, and the phylogeny of the group was reviewed. The degree of phylogenetic differences both between and among different coliform genera is indeed large, raising questions about their current definition. Using the database, current and new primer sets were tested for specificity and coverage both *in silico* and *in vitro*. The de novo primer set, LZ1, was found to be the most effective for qPCR. When compared directly with traditional culture-based methods and flow cytometry total cell counts, the LZ1 *lacZ* qPCR assay quantified *E. coli* in drinking water down to a concentration of 1.82 x 10^3^ cfu/100mL, equivalent to 360 cfu per qPCR reaction.

**Importance:** Coliform bacteria and E. coli are key indicators of faecal contamination in drinking water. Compliance with U.K., U.S.A., or European regulations requires that drinking water be free of these organisms. However, the culture-based methods used to quantify these bacteria have remained unchanged for over 100 years, relying on the culturability of the organisms, sampling only a small volume at a single time point, and lacking direct correlation to pathogenic microbes. There is a pressing need for a rapid, high-throughput analysis capable of detecting coliforms more representatively, without relying solely on culturability. By analysing larger volumes or more replicates, this qPCR assay could potentially be used to assess the risk of drinking water contamination over longer periods of time, offering a more comprehensive evaluation compared to traditional, culture-dependent methods.

## Introduction

### Coliform Bacteria

Since its discovery in 1892, *Escherichia coli* (1) has been an important indicator of faecal contamination in drinking water supplies, due to its presence in the enteric tracts of mammals. As microbiology developed in the early part of the 20^th^ century and other enteric bacteria were discovered, a group of bacteria, known as coliforms (and *E. coli* specifically) became and remain the primary bacterial parameters to assess the risk of enteric illness from drinking water. The definition of a coliform has changed over time. Traditionally they have been defined by their ability to be recovered by the methodology used to isolate them. Coliforms were first characterised by the production of gas from the fermentation of lactose and indole production, which is typical in *E. coli* species (2–4). However, in 1956 it was discovered that non-lactose fermenting species existed, so the classification of coliforms was changed to include genera like *Klebsiella, Enterobacter* and *Citrobacter* (5, 6). In 1996 the World Health Organisation (WHO) further expanded the definition to include an increasing number of variant species of coliforms, as ß -galactosidase activity was known to be a requisite for the group, but growth in the presence of bile salts and acid and gas formation within 24-48 hours from the fermentation of lactose was also required (7). In 1998 this was simplified to members of *Enterobacteriaceae* capable of producing ONPG (ortho-nitrophenol ß –galactosidase) (8). We now know that coliforms are not a strict phylogenetic group (9–12).

Coliform tests, alongside detection of protozoa like *Giardia* and *Cryptosporidium*, have driven public health improvements by limiting faecal contamination of drinking water supplies and therefore the spread of water-borne pathogens like cholera.

### Current Testing

Water utilities use a culture-based methodology for identification and quantification of coliform bacteria, that was developed in the mid 20^th^ century, and remains largely unchanged today. Testing is traditionally carried out by membrane filtration (MF) of drinking water where coliforms are trapped on a 0.25µm filter on top of a specific growth media and incubated at their optimum growth temperature (37°C). The growth medium contains ß-galactosidase and ß-glucorinodase (to distinguish *E. coli* from other coliform genera). The media contains a chromophore or fluorophore activated in the presence of these enzymes. Colour change of colony forming units (cfu) therefore determines a positive result. The regulatory standard is 0 colony forming units in 100 mL for coliforms and *E. coli*. There are several methods and proprietary media available, such as Multiple Tube Fermentation, and proprietary enzymatic systems like Colilert® or Colisure®, but all methods exploit the activity of ß-galactosidase and ß-glucorinodase.

More extensive confirmatory tests are needed alongside ß-galactosidase and glucorinodase activity. These may include further isolation of colonies and incubation on several media types at 37 and 44°C, then indole and oxidase tests. Isolated colonies may be further analysed using API (Analytical Profile Index) testing e.g. Biomerieux’s VITEK system® (13). Vitek uses the results of numerous biochemical tests contained within a sample cartridge, in an algorithm which predicts the most likely species of gram-negative bacteria. More modern confirmation techniques use matrix-assisted laser desorption/ionization time-of-flight mass spectrometry (MALDI-TOF MS) to analyse the ribosomal proteins of a bacterial cell to classify it. This has been proven to be a very effective method of classifying bacteria isolated using other methods (14).

### Limitations

There is an abundance of research from the past twenty years outlining the limitations of these methods. In general, culture-based testing is labour and time-intensive and in this case can take over 48 hours from sampling to result (where confirmatory tests are required). This means that water quality problems may have already become significant in a community before a utility can act. Culture-based testing is also known to only provide limited recovery of overall bacteria as >99% may be viable but not culturable in drinking water systems (Colwell et al., 1985; Oliver, 2005, Berney et al., 2008; Hammes et al., 2008; van der Kooij et al., 1995). There is evidence that MF techniques have an inability to recover stressed or injured coliform bacteria specifically (11), therefore current methods may further underrepresent the true concentration of these organisms in drinking water systems, particularly those containing residual biocides like chlorine.

All current enzymatic testing methods report a high number of false positives and negatives in coliform detection. Chromocult™, MI, Colilert™ and DC with BCIG agar have a recovery rate between 23-70% for non-*E. coli* coliforms (20, 21). The ISO-9308-1 method outlined by EU standards cannot distinguish *E. coli* from phenotypically similar organism *Klebsiella oxytoca*, which pose significantly lower health risks than *E. coli*. This method is unlikely to be able to quantify faecal coliforms (Fricker et al., 2008) and has also been demonstrated to fail to recover 15-16% of coliforms (Fricker et al., 2008). ISO and Colilert™ methods have been shown to demonstrate bias toward certain genera. Enterobacter genera such as *E. nimipressuralis, E. amnigenus, E. asburiae,* and *E. hormaechei* are commonly recovered as is *Serratia fonticola*. These species only ferment acid and produce gas after extended incubation. They are also unlikely to be faecal in origin (24). Additional confirmatory tests will take a minimum of a further 24 hours, further extending the response time to any water quality incidents. Furthermore, faster proprietary methods like Colisure™ and Colilert™ are susceptible from interference from turbidity, limiting their application where they might be most needed, i.e., during water quality incidents where both turbidity and bacteria may be elevated (25). No one method of coliform recovery is capable of recovery of all strains found in drinking water, whether of faecal or environmental water.

Despite these issues, water regulators rely on the absence of coliforms (0 cfu 100ml^-1^) in routine samples to demonstrate the safety of the drinking water supply. Detections are extremely low in frequency (<0.1% samples) and magnitude (∼1 cfu 100ml^-1^) but remain at persistent low-levels for most water utilities (26). Therefore, there is a need for a more timely, accurate and meaningful analysis to identify and quantify coliform bacteria in drinking water.

### Quantitative Polymerase Chain Reaction (qPCR)

Quantitative polymerase chain reaction (qPCR) is used in a wide variety of industries to identify species or genera-specific gene sequences using a fluorescent probe. qPCR analysis is much faster than culture testing (e.g. COVID-19 testing from wastewater using qPCR provided results in <24hours, ref). qPCR also has the advantage over culture methodologies in that it can be highly specific to its target over a dynamic range of target concentration. qPCR has been employed in the water industry to track sources of contamination, primarily to track sources and strains of the pathogen *Bacteroides*. *Bacteroides* strains are very specific to their hosts organisms and can therefore be used to identify the source of contamination of a water supply (27–29). In a similar manner qPCR could be used to quantify indicators like coliforms and *E. coli*. However molecular analyses have yet to be widely adopted by water utilities, as regulators still direct utilities to carry out culture testing (30).

There are several studies using qPCR and other molecular tools, like 16S rRNA amplicon sequencing to characterise and quantify microbial communities in drinking water (31–37). These have revealed critical factors that determine the abundance and type of bacteria within drinking water distributions (DWDS), including the quality of the source and filtration in seeding communities downstream. However, more research is needed to better understand the role of coliforms and *E. coli* in the drinking water microbiome, so that water utilities can reduce coliform detections and improve the quality of their supplies (38, 39). A robust molecular method for quantifying and identifying coliform bacteria in drinking water is required, to support culture-based microbial testing. This could be used to identify areas of risk for coliform or other pathogen detections across a typical treatment system and investigate correlations between coliform bacteria and pathogens within drinking water systems in general. By understanding these factors more proactive risk management could be adopted, reducing the cost and time spent on failure investigations, and reduce the risk of public health issues.

To date several studies have attempted to develop molecular techniques to directly quantify coliforms in drinking water. This is extremely challenging, as they are not a phylogenetically or functionally related group. Therefore, most PCR studies target the gene encoding ß-galactosidase (*lacZ*), as ß -galactosidase activity is the one requirement for classification using culturing. The first primer set, by Bej et al., in 1990 was updated in 1991 (1675F/2025R) using an expanded sequence database. Several studies have used these to target coliforms in the environment (40–47). However, these primers have low specificity and recover non-coliform organisms including *Salmonella* or *Shigella*, who also have *lacZ*. These primers also failed to target all known coliform genera (9). Two more recently developed primer sets report improved specificity: *lacZ*3F/R and *lacZ*3153f/3995R (9, 48). These have better coverage than 1657f/2025R but still amplify *Shigella* and *Salmonella* species (9, 48). Specificity testing of these primers was carried out using a small group of known coliforms, and all were developed using a small database of sequences (∼n=100). None of these have been used in qPCR assays. Further expansion, testing and application of *lacZ* primers are required, with the aim to increase their ability to recover a wider range of coliforms and use them quantitatively.

The aim of this work was to develop a robust qPCR assay for coliform bacteria. To do this, first a database and phylogeny of all known *lacZ* sequences was created. This was then used to evaluate *in silico* coverage and specificity of current *lacZ* primers and generate *de novo* primers. The best performing primers were tested *in vitro* against a test panel of typical coliform and non-coliform organisms isolated from drinking water. Lastly the successful primers were optimised for use in qPCR. The sensitivity of the qPCR assay was also tested against traditional MF MLGA culture testing and flow cytometry to identify the limits of detection.

## Materials and Methods

### Database collation

A database of 1427 putative *lacZ* sequences was generated using a confirmed *lacZ* sequence (*Klebsiella pneumoniae spp*) from the UniProt® website to retrieve sequences. A blastn search was carried out for highly similar sequences in NCBI®, limited to the *Enterobacteriaceae* phylum. This was pruned to include sequences of similar length (∼3000 bp) and aligned using MAFFT (Multiple Alignment using Fast Fourier Transform) to generate an aligned FASTA file (49) of 1292 *lacZ* sequences. MAFFT alignments are particularly good for large volumes of sequences, as they use obvious regions of homology to build the alignment, allowing faster processing without compromising on accuracy. The database and aligned fasta file are available in the Supplementary information (Supplementary file 2). *LacZ* primers were generated and tested using the program PrimerProspector with the aim of developing a robust target for qPCR in terms of product size and projected analysis conditions. Primer Prospector has been successfully applied to testing primers in several environments including soil and surface water, making it ideal for this analysis (50, 51).

### Phylogeny

The phylogeny of the *lacZ* database was explored using MRBAYES, a program using Bayesian inference to generate the most likely *lacZ* tree from over 100,000 trees generated from the MAFFT alignment (52). To compare *lacZ* phylogeny to a reference gene (16S rRNA), whole genome records for all individuals in the database were searched using universal primer set 27f/1492r in PrimerProspector to generate 16S rRNA sequences. 100 amplicons for 16S rRNA were identified, aligned using MAFFT and passed to MRBAYES. The trees generated from both analyses were modelled using BUCKy, to find a concordant phylogeny. BUCKy uses a Dirichlet process, does not infer the source of discordance, and does not require sequences to be assigned to a species, unlike other gene models. The concordance is based on the clustering of genes with the same topology (53). The individual concordant trees from MRBAYES were also analysed using PHYANOVA to identify whether speciation or inclusion as a coliform explained the variance in the distance between individuals in the database. Phylogenetic trees were visualised using the International Tree of Life (iTOL) program (54).

### In silico testing

Potential *de novo* primers were identified using this database in PrimerProspector software’s generate_de_novo_primers.py module. This generated a file containing all database hits and their associated score and sequence. Scores generated by PrimerProspector weigh different parameters in a combined score (Weighted score = non-3’ mismatches * 0.40 + 3’ mismatches * 1.00 + non-3’ gaps * 1.00 + 3’ gaps * 3.00. An additional 3.00 penalty is assigned if the final 3’ base mismatches), where the lowest scoring primer provides the best potential coverage (55). The lowest scoring primers were analysed alongside 3 existing primer sets (9, 40, 41, 48) against the unaligned *lacZ* database in PrimerProspector and amplicons generated for each forward primer with each reverse. To test the variation in the distance between individual sequences within the database by their species as well as whether they were designated a coliform, analysis of variance was tested (PHYANOVA) using R’s Vegan package (56). This was used to identify the primer set that could best distinguish between coliform and non-coliform members of *Enterobacteriaceae*, where the highest amount of variation between coliforms and non-coliforms is the most discriminatory primer set. This was required to ensure false positives were not identified in the analysis. The best performing primer sets were then optimised using PrimerProspector to identify whether the inclusion of any redundant base pairs improved the number of amplicons generated *in silico*. The sets were then re-scored to test optimisation. Base pairs were altered within the primer if more than 5% of the database was not covered by the current base.

### Test Panel and PCR

The top three primers were taken forward for *in vitro* testing using a test panel of organisms. Test panel organisms were a mixture of laboratory strains cultured in 50 mL LB and grown overnight at 37 °C and environmental organisms isolated from untreated water samples. Untreated water was filtered on 0.45µm membrane filters and grown on MacConkey Lactose Gentamycin Agar (MLGA (Oxoid®) prepared following manufacturer’s instructions. Samples were incubated at 37°C for 24 hours. A selection of single yellow (Coliform) and green (*E. coli*) colonies isolated from the MLGA plates were then inoculated into Phosphate Buffered Saline (PBS). DNA from all organisms was extracted using Qiagen DNeasy PowerWater Kit following manufacturer’s instructions and eluted in 100 µL elution buffer. DNA concentrations were measured using Qubit® following manufacturer’s instructions and samples diluted with Ultra-Pure (UP) water to 2ng µL^-1^ DNA.

For end point PCR cycle Bioline’s MyTaq™ PCR kit was used in 25µL sample volumes, with a final primer concentration of 0.4 µM. For existing primers, PCR conditions were as outlined in the original research, conditions can be found in Supplementary Information (Supp Table 3). For *de novo* primers, the optimal annealing temperature was found by running an initial temperature gradient PCR. Final cycle conditions for LZ1 *de novo* primers were: 95°C for an initial 5 min then 30 cycles of 95 °C for 30 seconds, 50°C for 30s; 72 °C for 1 min. For qPCR, reactions were carried out using a BioRad® thermocycler and Quantitect® SYBR Green qPCR Kit in 20µL volumes. 2µL of template from each sample was added, and primers to a final concentration of 0.3 µM. Cycle conditions for qPCR were as end-point reaction except there were 40 cycles. A final melt-curve analysis was carried out at from 65 to 95 °C. Standard curve was constructed from 10^3^ to 10^8^ using a synthetic DNA amplicon. The sequence used for the block was the consensus sequence from amplicons generated in PrimerProspector and aligned in MAFFT for LZ1. This is available in Supplementary information.

### Quantification Testing

To test the detection limits of the best performing primer set, an experiment was set up to compare the enumeration of the qPCR assay compared to a typical MLGA culture test. *E. coli* K12 was inoculated from a single colony into LB and incubated overnight at 37°C. The culture was then enumerated using flow cytometry to measure cell counts. For this, 100µl of culture was diluted 1:100 with 1x SYBR green and incubated at 30 °C for 15 minutes. 50µL of sample was tested by the BD Accuri Flow Cytometer using the standard gating protocol for drinking water (57). Cells were then serially diluted in PBS and inoculated in triplicate into a background of drinking water or pure water representing a log dilution series of *E. coli* from 10^8^ to 10^1^ cells per 100ml, plus a control within no added *E coli*. Coliforms within each water sample were subsequently enumerated by 1) filtering 100 mL onto 0.45 µm and incubating the filter on MLGA agar for 24hrs at 37C; 2) filtering 100 mL onto 0.45 µm, the filter was then frozen at -20 °C until subsequent DNA extraction using Qiagen DNeasy PowerWater Kit and eluted in 100 µL. DNA was further concentrated using speed vacuum to 10 µL of DNA (representing 10^X^ cfu 100mL). 2 µL of this DNA was used for qPCR quantification of coliforms using primer set LZ1. qPCR testing conditions were as before: 95°C for an initial 5 min then 40 cycles of 95 °C for 30 seconds, 50°C for 30s; 72 °C for 1 min. A final melt-curve analysis was carried out at 95 °C. Standard curves were constructed as above. Finally, FCM quantification within the water dilution was carried out using the same method as above.

## Results

### Phylogeny of *lacZ* and 16S rRNA

A phylogeny of *lacZ* sequences is outlined in Figure 2. There are numerous distinct groups of non-coliform bacteria, with *Vibrio, Aeromonas and Salmonella* genera forming distinct clades. *Aeromonas* was closest in distance to coliform genera and more distinct from the other non-coliform groups. *Salmonella* and *Vibrio* were more closely related to each other. Of the coliform sequences, *Enterobacter* species spread widely across the tree, occurring on four distinct branches, one of which contained non-coliform *Vibrio* sequences. Many of the other coliform sequences do cluster with their species, however large numbers of the sequences were too closely related to distinguish distinct clusters. *Klebsiella* species were the most genetically distant to the other coliform genera. The phylogeny was generated using sequences of similar length from the initial database, giving 1292 putative *lacZ* sequences. Overall, genetic distance between sequences was low, with most branch lengths <0.05.

**Figure 1:**
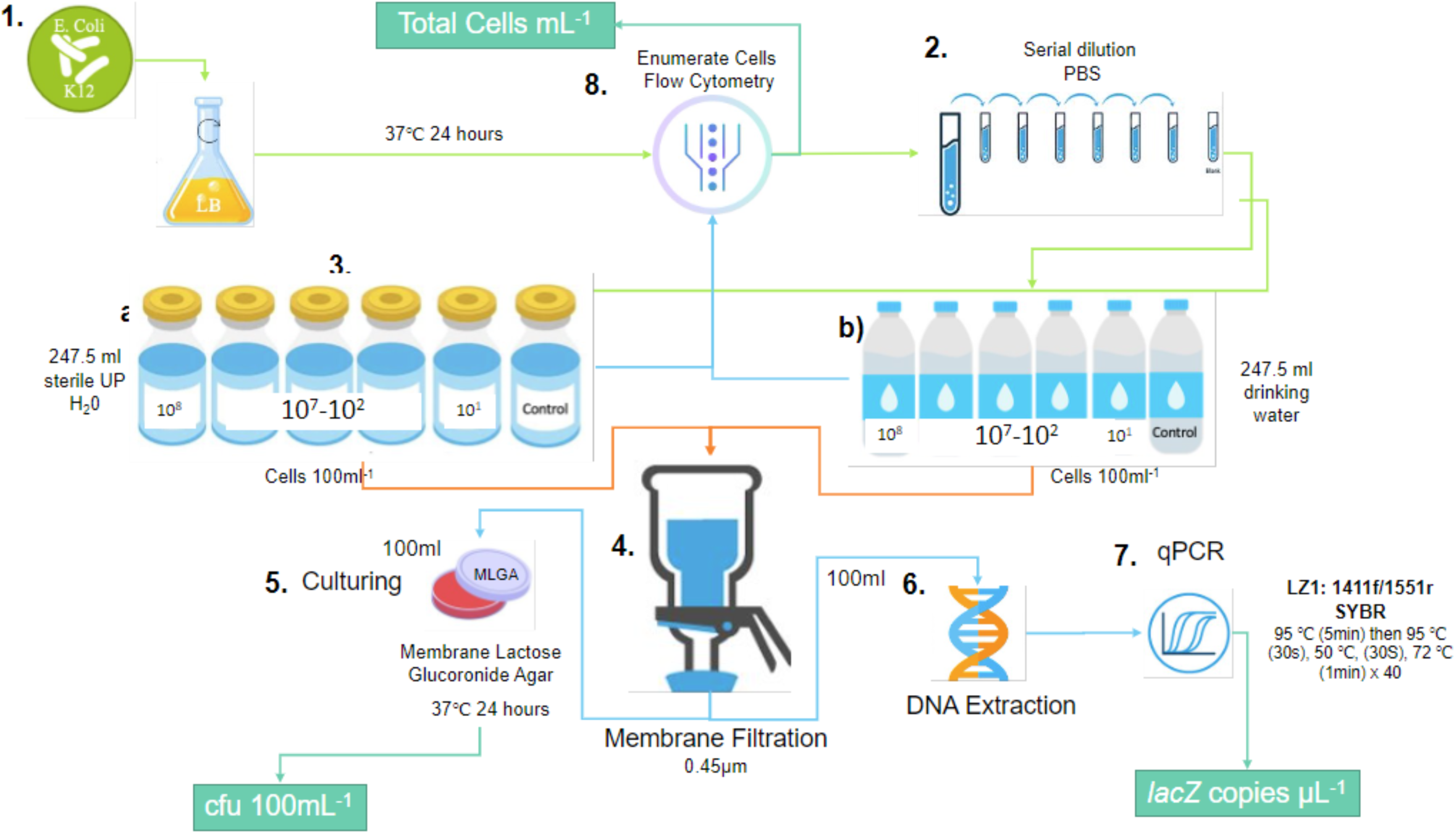
Overview of in vivo qPCR testing of LZ1. 1. *E. coli* K12 was inoculated from a single colony into in LB and grown overnight at 37 °C. 2. Total cells mL^-1^ were then enumerated using flow cytometry, and a serial dilution of the culture made in PBS. 3. Cells were added to a) Ultra-pure water and b) tap water in a range of concentrations (10^8^ to 10^1)^ per 100mL. 4. 100mL of each Sample was filtered onto 0.45um membrane filters and cultured on MLGA (5.). 6. A second 100mL was filtered and DNA was extracted. 7. qPCR testing of DNA extractions carried out using LZ1. 8. Flow cytometry of spiked samples for total cell count.

**Figure 2:**
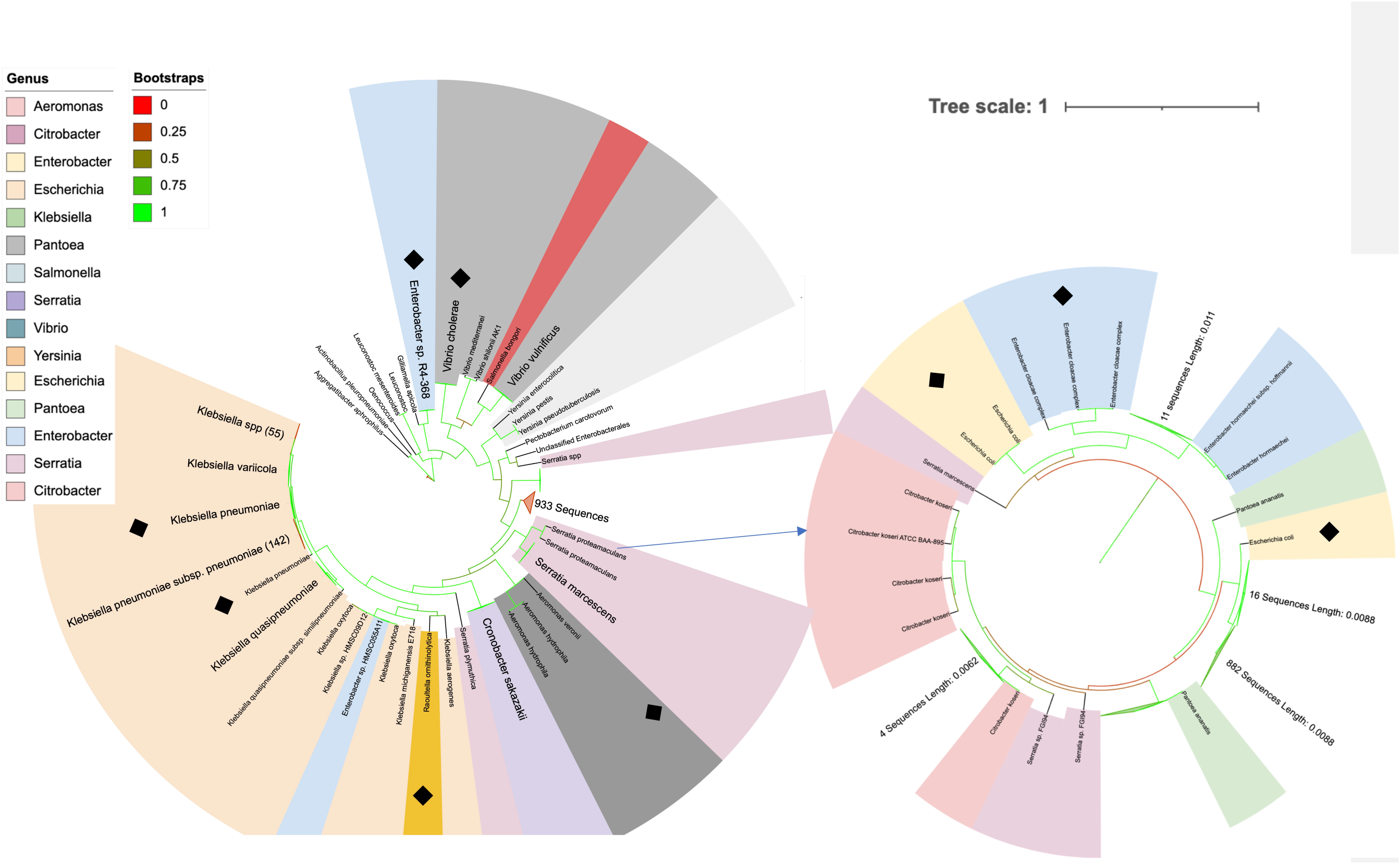
Phylogeny of /acZ sequences aligned using MAFFT, n=1292. a) Branches with Bootstrap values >0.55 are collapsed. b) Phylogeny of collapsed clade with 933 sequences (circled on a)). Branch lengths <0.05 are collapsed. Black diamonds denote that these species were tested as part of this study.

When compared to the concordant phylogeny of both 16S rRNA and *lacZ* sequences in Figure 3, this is distinctly different. This phylogeny contains significantly less sequences (n=100) due to the requirement for both full-length sequences in NCBI (**Error! Reference source not found.**). There are distinct groups loosely associated with species type. As in the *lacZ* phylogeny *Klebsiella pneumoniae* species are the most distant from the other coliform genera. There are several clusters of *E. coli*. though it is noted there are unclassified *Enterobacteriaceae* between these groups, which could be *E. coli*. The large number of unclassified sequences to genus or species level in the database further highlights the difficulties in classifying coliform bacteria. Many individual species do not cluster together in the coliform group in the concordant phylogeny. *Salmonella enterica* species were not closely related to any of the *E. coli* groups, but *Shigella* species were. This further highlights the genotypic similarities *Shigella* and *E. coli* share, often more closely than other species of *E. coli* (58).

**Figure 3:**
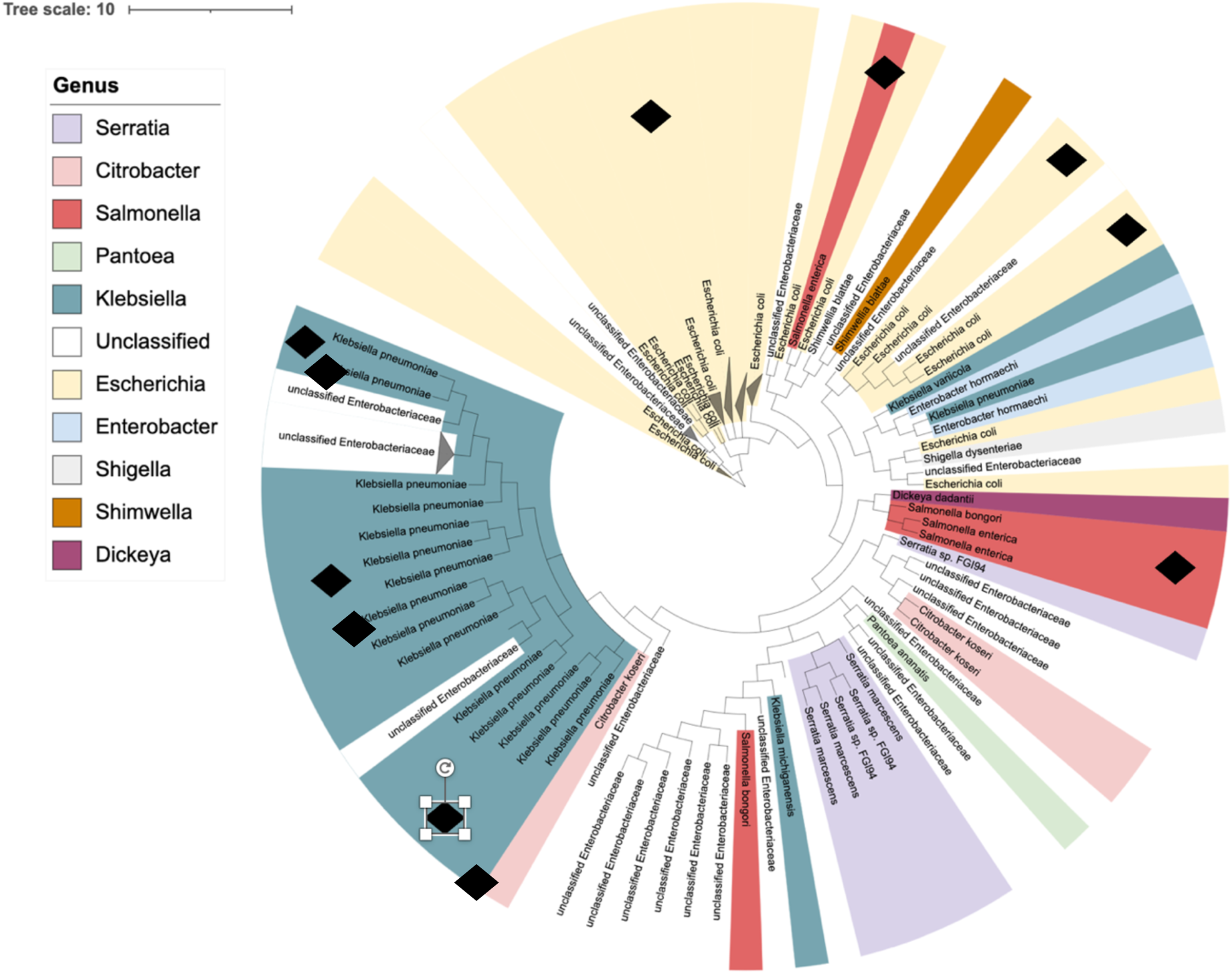
Output of BUCKy software, showing most likely concordant phylogeny of /acZ and 16S rRNA gene trees. Node values show concordance factors for the grouping. Black diamonds denote that the species was tested as part of this study.

When compared to the concordant phylogeny of both 16S rRNA and *lacZ* sequences in Figure 3, this is distinctly different. This phylogeny contains significantly less sequences (n=100) due to the requirement for both full-length sequences in NCBI (**Error! Reference source not found.**). There are distinct groups loosely associated with species type. As in the *lacZ* phylogeny *Klebsiella pneumoniae* species are the most distant from the other coliform genera. There are several clusters of *E. coli*. though it is noted there are unclassified *Enterobacteriaceae* between these groups, which could be *E. coli*. The large number of unclassified sequences to genus or species level in the database further highlights the difficulties in classifying coliform bacteria. Many individual species do not cluster together in the coliform group in the concordant phylogeny. *Salmonella enterica* species were not closely related to any of the *E. coli* groups, but *Shigella* species were. This further highlights the genotypic similarities *Shigella* and *E. coli* share, often more closely than other species of *E. coli* (58).

PHYANOVA analysis of the 16S rRNA, *lacZ* and concordant phylogenies is shown in Figure 2. The variance in phylogenetic distance between coliforms and non-coliforms (group) was not significant for any of the trees, with the concordant phylogeny having the least, <1%. *lacZ* had more variation explained by grouping at 12%. This suggests *lacZ* is more variable between non-coliforms and coliforms than the 16S rRNA gene. The amount of variation explained by species was also higher within the *lacZ* gene at 26%. A study of sequences in *Enterobacteriaceae* suggested that the 16S rRNA gene was not an ideal marker for a phylogeny of the family, due to the multiple copies and the high level of conservation (10). Zhang et al (2015) produced a 16S rRNA phylogeny of the coliform group which successfully distinguished coliforms into 32 distinct groups based on 16S rRNA, but required a combination of *lacZ, uidA* and 16S rRNA targets to successfully target coliforms. The number of sequences recovered in this study was very small (n=100). A larger study of both *lacZ* and 16S rRNA sequences recovered from wild types from various environments is required to understand the lineage of enteric bacteria more fully, and their role in disease. This will also help identify potential markers that can be used to identify the group overall.

**Figure 2:**
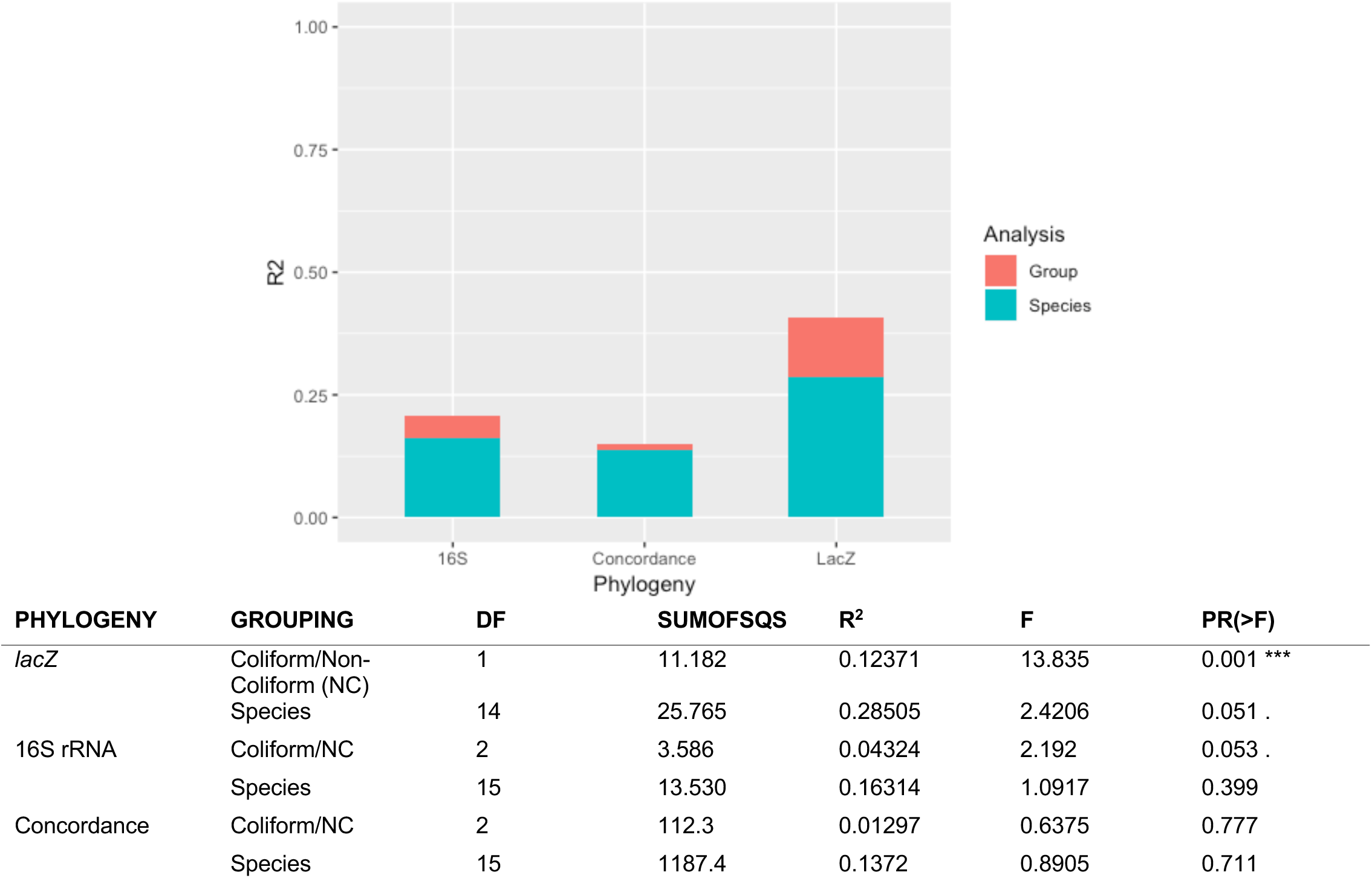
PHYANOVA results of comparison of genetic distance between coliform/non-coliform (Group) and by Species, for 100 sequences in the lacZ database with a corresponding 16S rRNA sequence. R2 values are given for variation in the lacZ and 16S rRNA phylogenies as well as a Concordant phylogeny generated by BUCKy.

### In silico results

PrimerProspector generated 2348 primers from the initial *Klebsiella oxytoca* sequence. These were tested against the 1292 *lacZ* sequences in the database. The database was further subdivided into coliform (951) and non-coliform (214) and unclassified *lacZ* sequences to identify primers which best distinguish between these groups. Each forward and reverse primer was tested in isolation to generate the number of database hits. Table 1 outlines the best performing existing and *de novo* primers identified, alongside their PrimerProspector score. Details of the sequence database and the primers tested can be found in supplementary information (Supplementary table 4). 1411F and 1551R had the best mean scores of the primers generated by the program, 0.52 and 0.43. Of the existing primers, *lacZ*3153f/3995r (9) scored most highly: <0.65 with the highest number of perfect scoring database hits (68%). The poorest scoring primers were 1675F and 2025R, with a mean score >2 and 33% and 57% of hits scoring 0. The best performing existing primers (3153f/3995r), have similar proportion of redundant bases (2–3) to the *de novo* primers except for RP6. This primer is unlikely to be viable, as it has 7 redundant bases. It was included as an outgroup to test bias against primers with high levels of redundancy. This primer scored 0.788 but had 54% of hits with a perfect score.

**Table 1:**
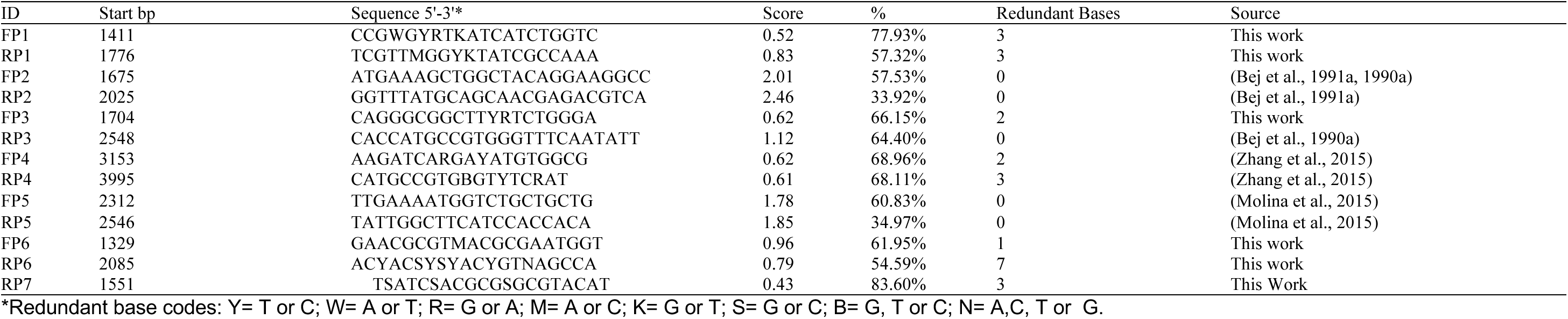
Details of *lacZ* primer sets tested using PrimerProspector. Start bp is given as described in prior work. Score is mean weighted score= non-3’ mismatches * 0.40 + 3’ mismatches * 1.00 + non-3’ gaps * 1.00 + 3’ gaps * 3.00. An additional 3.00 penalty is assigned if the final 3’ base mismatches. % is number of database hits scoring 0 (n=1427).

**Table 2:**
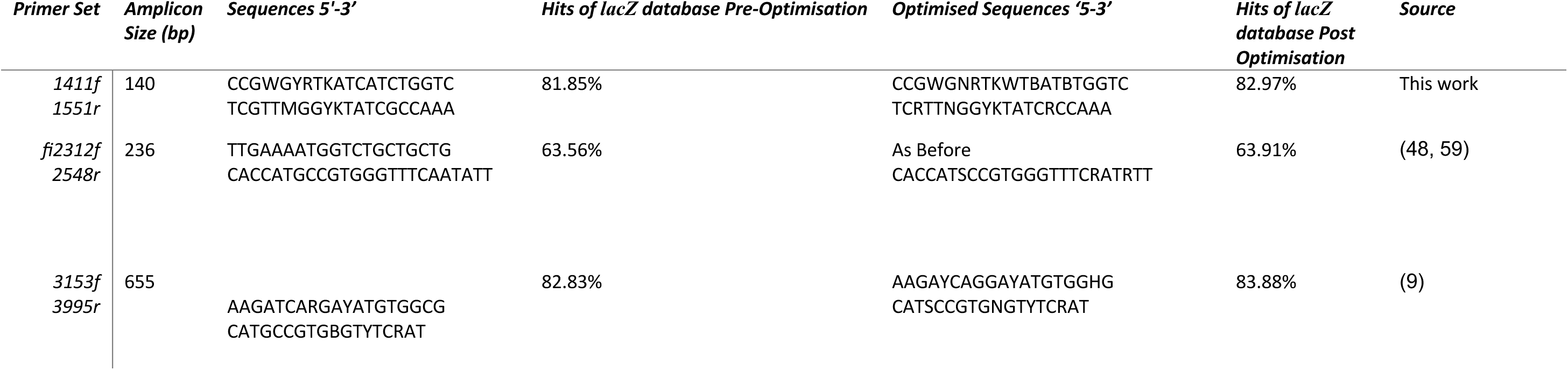
Original and optimised primer sets with significant coverage across the *lacZ* database. Also showing the amplicon size generated and the percentage of amplicons returned /1427 sequences (% Hits) and the source of the primer.

Primers were then tested in pairs (Figure 3). Amplicons from each primer set were aligned and distance matrices calculated between individual sequences. As with the full-length sequence phylogenies, Analysis of variance was carried out on each distance matrix to test whether there were significant differences in the distance matrices between species or between coliforms and non-coliforms (Group in Figure 4). No primer set generated amplicons with R^2^ > 20%, suggesting coliforms and non-coliform sequences do not have a significant difference in terms of distance (where R^2^ >0.2 p=<0.05). Speciation explained more variation with >99% variation in the amplicons of the primer set 1368f/1776r (FP1_RP1) explained by their species. The primers with the most hits across the database 1411F, 3153F, 3995R and 1551R had the lowest R values.

**Figure 3:**
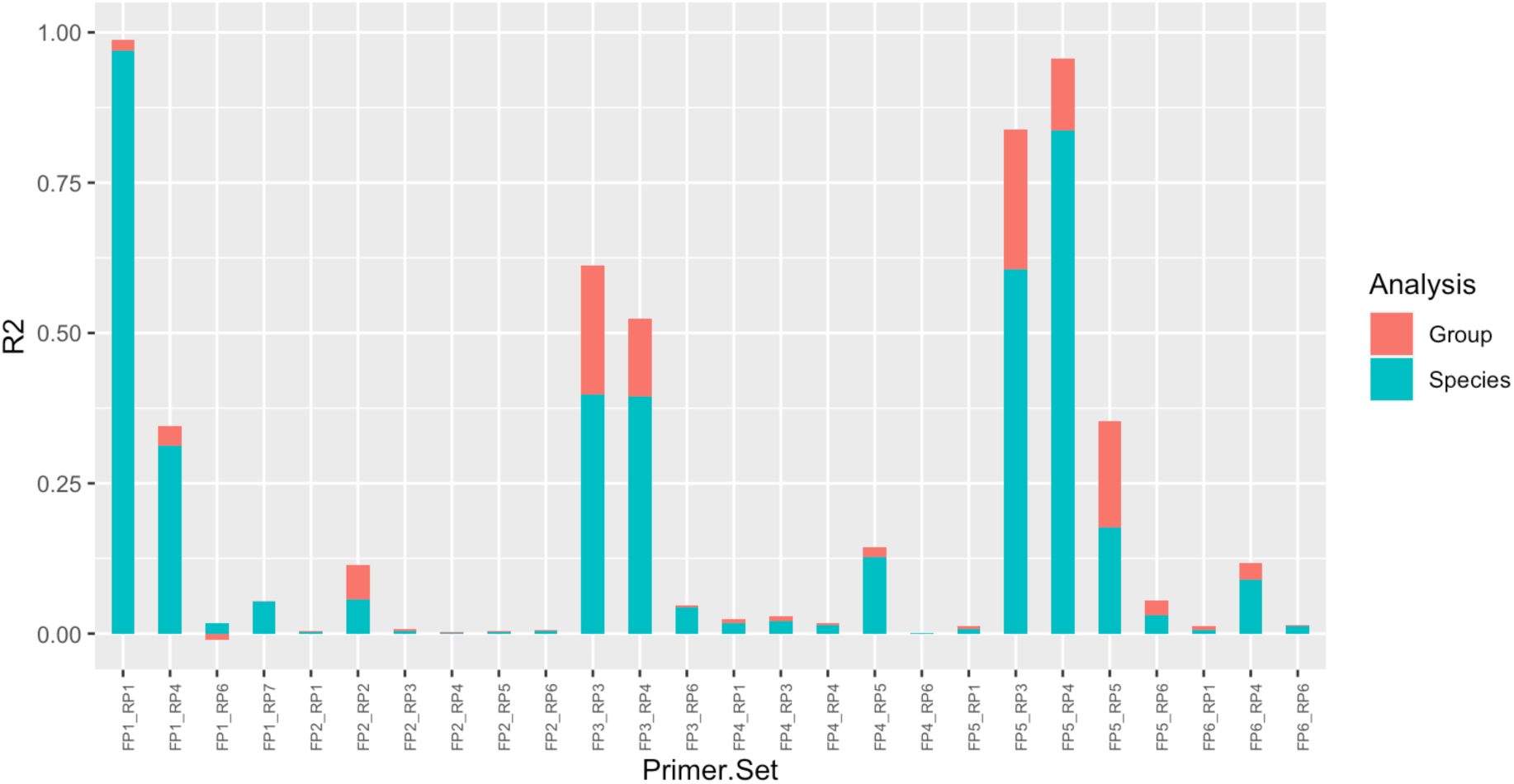
Results of PHYANOVA test on all amplicons generated from primer sets outlined in table 1, where amplicons generated were >3. The R2 value describes the proportion of variation explained by the sequences being from a coliform (Group) and amount of variation explained by Species groupings (blue). FP1_RP1, FP5_RP3 and FP5_RP4 all had the most variation in the amplicons by Group and by Species, indicating that these are good primer sets to distinguish non-coliform Enterobacteriaceae (all p=0.01).

**Figure 4:**
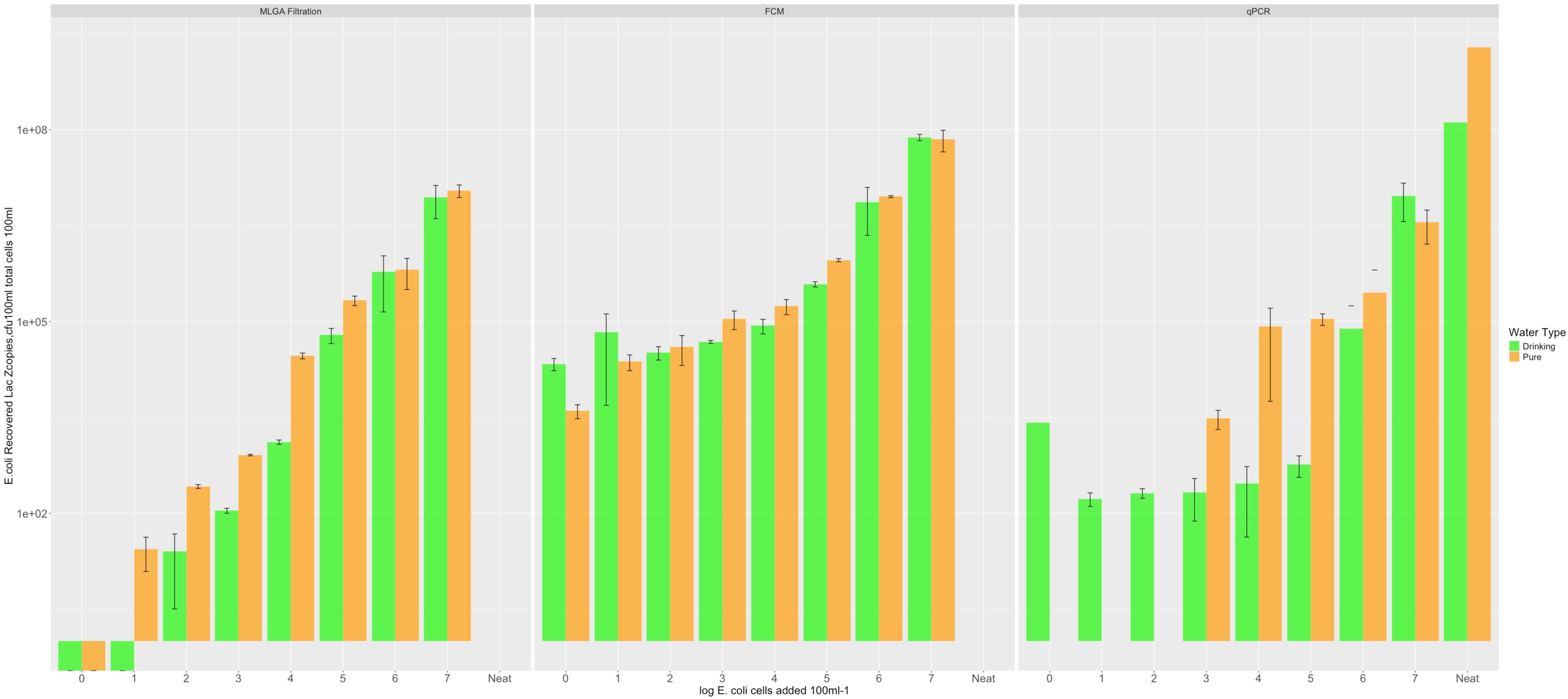
Results of sensitivity testing of *lacZ* primers LZ1. Shared x axis shows the theoretical log concentration of *E. coli* 100mL^-1^. Top: Mean flow cytometry results in total cells mL^-1^, middle, culture testing in cfu 100mL^-1^ and bottom qPCR results in *lacZ* copies µL^-1^. Error bars show standard error on the mean and are based on triplicate analysis. Drinking water samples are shown in green, pure water samples in orange. Neat is extraction of DNA from 100mL undiluted culture.

Based on these results, three primer sets were selected for further testing: 1411f/1551r (LZ1, from this study); 2312f/2548r (LZ2, Bej et al., 1991a; Molina et al., 2015) and 3153f/3995r (LZ3, Zhang et al., 2015). LZ1 and LZ3 scored most highly on the number of database hits- >82%. LZ3 generated large amplicons (655 bp), which makes it less efficient for qPCR, particularly without an available probe site. LZ3 were compared in a previous study, where they were demonstrated to have an improved performance compared to those developed by Bej et al (Bej et al., 1990; Zhang et al., 2015), therefore, these were included for further testing despite their low R^2^ value. Primer 2312f had no change to primer sequence pre and post optimisation, and although scored lower in terms of database hits, had no redundant bases. This was interesting considering it overlapped primer 3153f. All primers had slight increases to their percentage hits post optimisation, but these were only around 1%. Therefore, original primer sequences were used (Table 3).

**Table 3:**
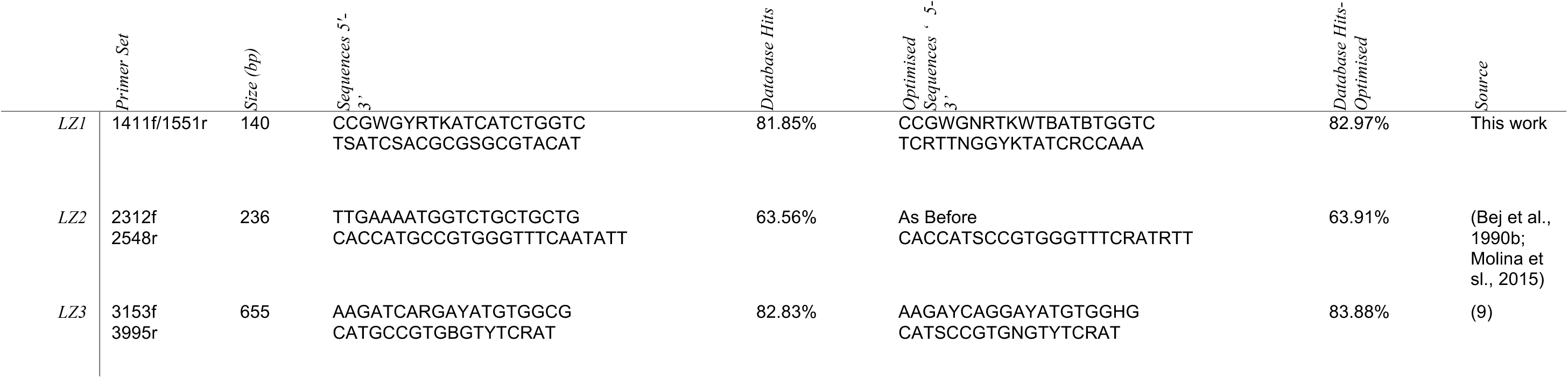
The 3 best performing primer sets in silico. Sequences were scored optimised using PrimerProspector software against the *lacZ* database. LZ1 was a de novo set identified in this work. LZ2 and 3 were primers from existing research.

### Test Panel

The three primer sets were tested against a range of coliform and non-coliform enteric bacteria (isolated from drinking water sources) (Table 4). LZ3 amplified the most coliform organisms, with a positive result for all organisms tested. However, it also amplified the most non-coliform organisms. LZ1 had the next highest percent recovery, 87.5%, however it failed to give a positive result for a laboratory strain of *Enterobacter aerogenes*, although it did amplify other *Enterobacter species*. LZ1 also had one positive for a non-coliform organism, *Aeromonas jaundii*. This species had an *in vitro* positive result for all three primer sets, and in addition appeared as a yellow colony when cultured on MLGA (indicative of being a coliform). *Aeromonas* species were also recovered by the primers *in silico* and were more closely related to coliforms than other typical non-coliform *Enterobacteriaceae* in the phylogenies (Figures 2 and 3). LZ2 was the poorest performing set, failing to positively identify a species of *E. coli*, *Hafnia alvei,* and *Enterobacter aerogenes*.

**Table 4:**
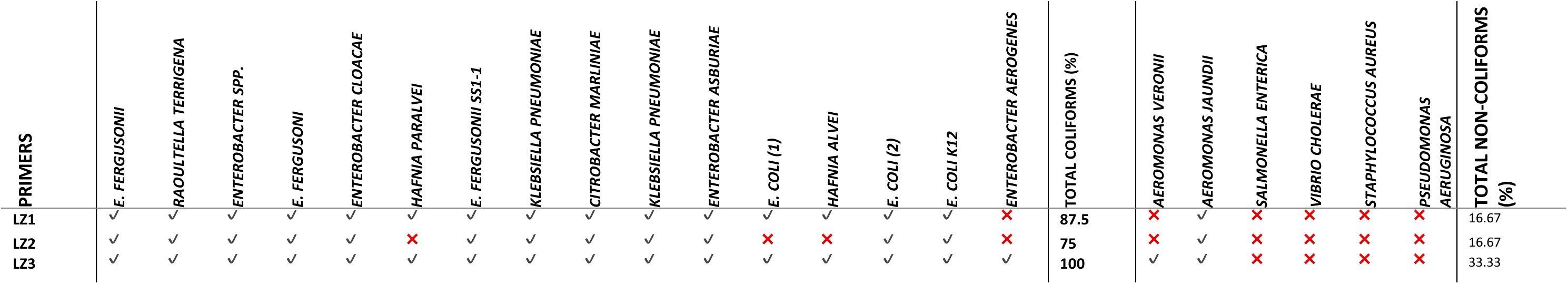
Test panel results for *lacZ* primers LZ1, LZ2 and LZ3. Tick denotes a positive result from end point PCR. Organisms isolated and cultured from the environment were identified from 16S rRNA amplicon sequencing. The percentage of coliform and non-coliform organisms recovered by each primer set is indicated.

### qPCR Sensitivity Testing

When tested using known concentrations of *E. coli* the qPCR results were mixed for both pure and drinking water samples (Figure 4). All MLGA filtration results were within the expected cfu 100 mL^-1^ ranges, except in drinking water, where there were no colonies at 1 x 10^1^ *E. coli* 100 mL^-1^. The flow cytometry total cell results followed a similar pattern. The blank samples with drinking water having an average of 216 cells mL^-1^ (21600 cells 100mL^-1^). compared to 26 cells mL^-1^ in the pure water. Drinking water and pure water samples were much closer in concentration to the MLGA testing, with drinking water being slightly lower, except at 1 x 10^2^ *E. coli* added. The linear regression of actual versus expected plate counts was within expected tolerances for Pure water samples, R^2^=0.96, and slightly lower in Drinking water samples R^2^=0.93 (Figure 6 a)).

**Figure 5:**
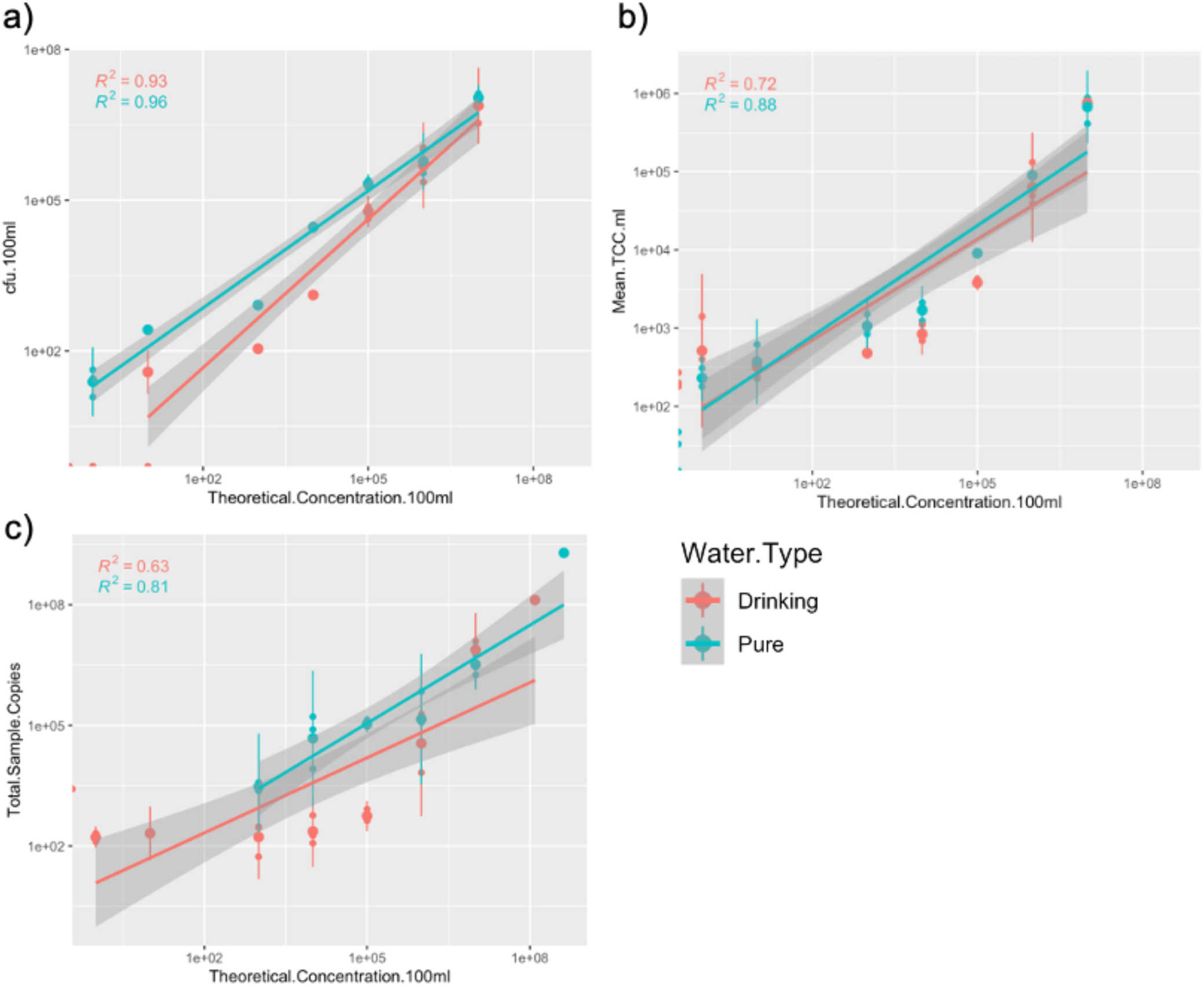
a) Linear regression analysis of expected versus actual results for a) MLGA Filtration cfu 100mL^-1^, b) Flow Cytometry total cells mL^-1^ and c) LZ1 qPCR total sample copies, for all drinking water samples (red) and ultra-pure water samples (green).

**Figure 6:**
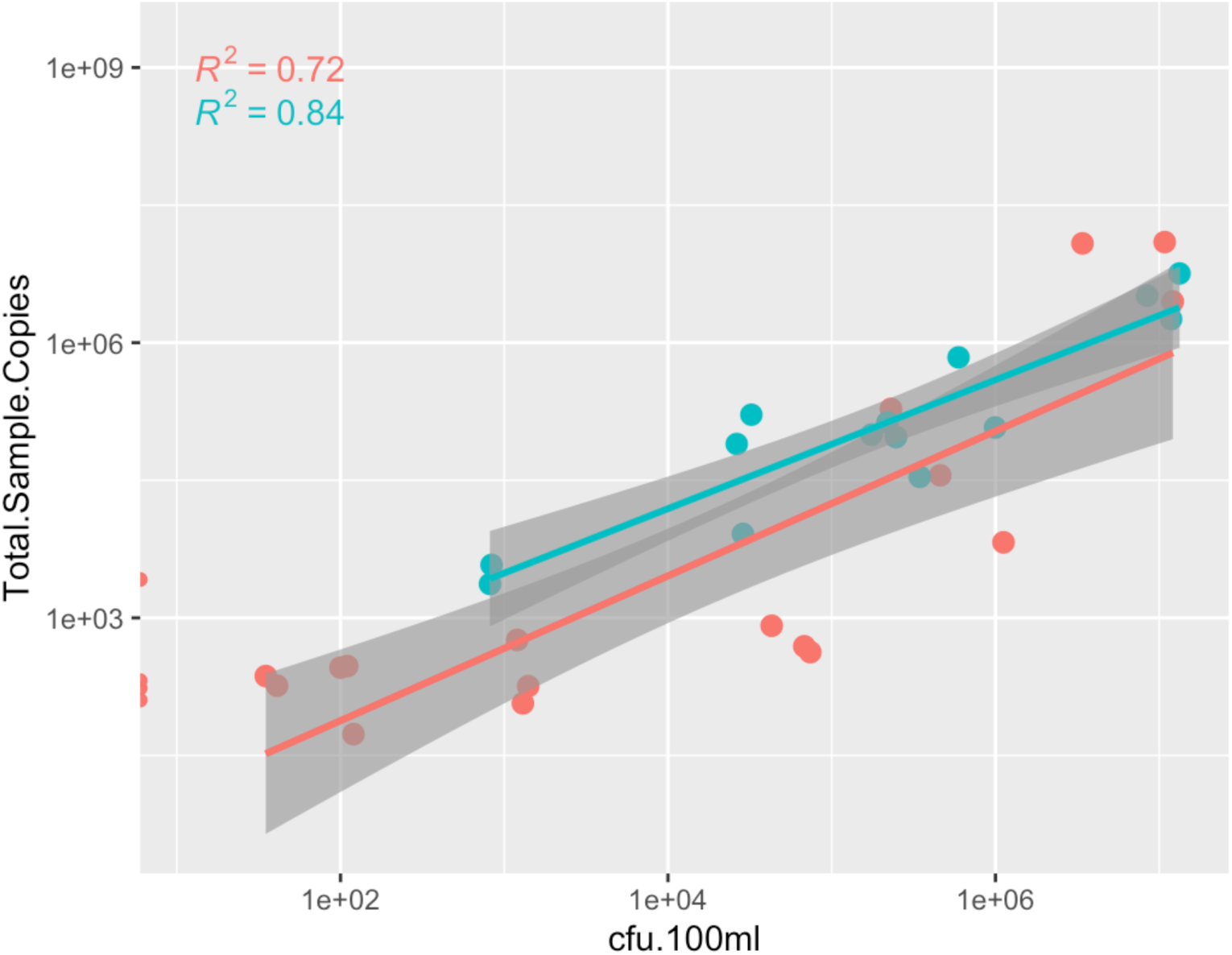
Linear regression of actual qPCR and MLGA filtration results. For all drinking water samples (red) and ultra-pure water samples (green).

The drinking water samples had a higher amount of background cells compared to the pure water samples, in the 0-1 log cells added range. But there was a much higher level of error associated with samples with <3 log cells added. The actual versus expected for total cell counts was not as strongly correlated as with the plate counts, but still strong R^2^=0.88 and R^2^=0.72 for pure and drinking water sample results respectively.

The qPCR results of the neat *E. coli* culture, for the drinking water samples had a total of 10.82 log copies compared to that prepared for the pure water which had 8.17 log copies, although MLGA cfu 100mL^-1^ were equivalent. These were prepared on different days from different cultures. The average blank control (no *E. coli*) for the drinking sample was 2.32 log copies, whereas there was no signal detected from the pure water samples. NTC samples were undetected. Standard curve had 89.7% efficiency, a slope of -3.595 and a y intercept of 36. 67 cycles (R^2^ = 0.98). The low concentrations (1-2 log *E. coli* 100 mL ^-1^) also had no signal detected for pure water samples, but a total of 1.68 and 2.07 log copies in the drinking water samples. Above 3 logs of *E. coli* added the pure water samples had 3.06 log copies compared to 815 cfu 100mL^-1^ on MLGA, following a similar pattern up to 7 logs *E. coli*. Drinking water samples had higher log copies than pure water in samples with log 3-6 *E. coli* added (5.78-9.13 total copies). This is around 2 logs higher than the pure water samples. The actual versus expected values for the qPCR were the lowest of the 3 methods, R^2^= 0.81 and R^2^=0.63. However, when directly comparing the actual culture testing results and actual qPCR log copies (Figure 7) samples have a higher correlation to culture testing, pure water, R^2^=0.84 and drinking water R^2^=0.72. Drinking water samples could have lower correlation in the qPCR tests due to signals in the blank and low concentration samples.

**Figure 7:**
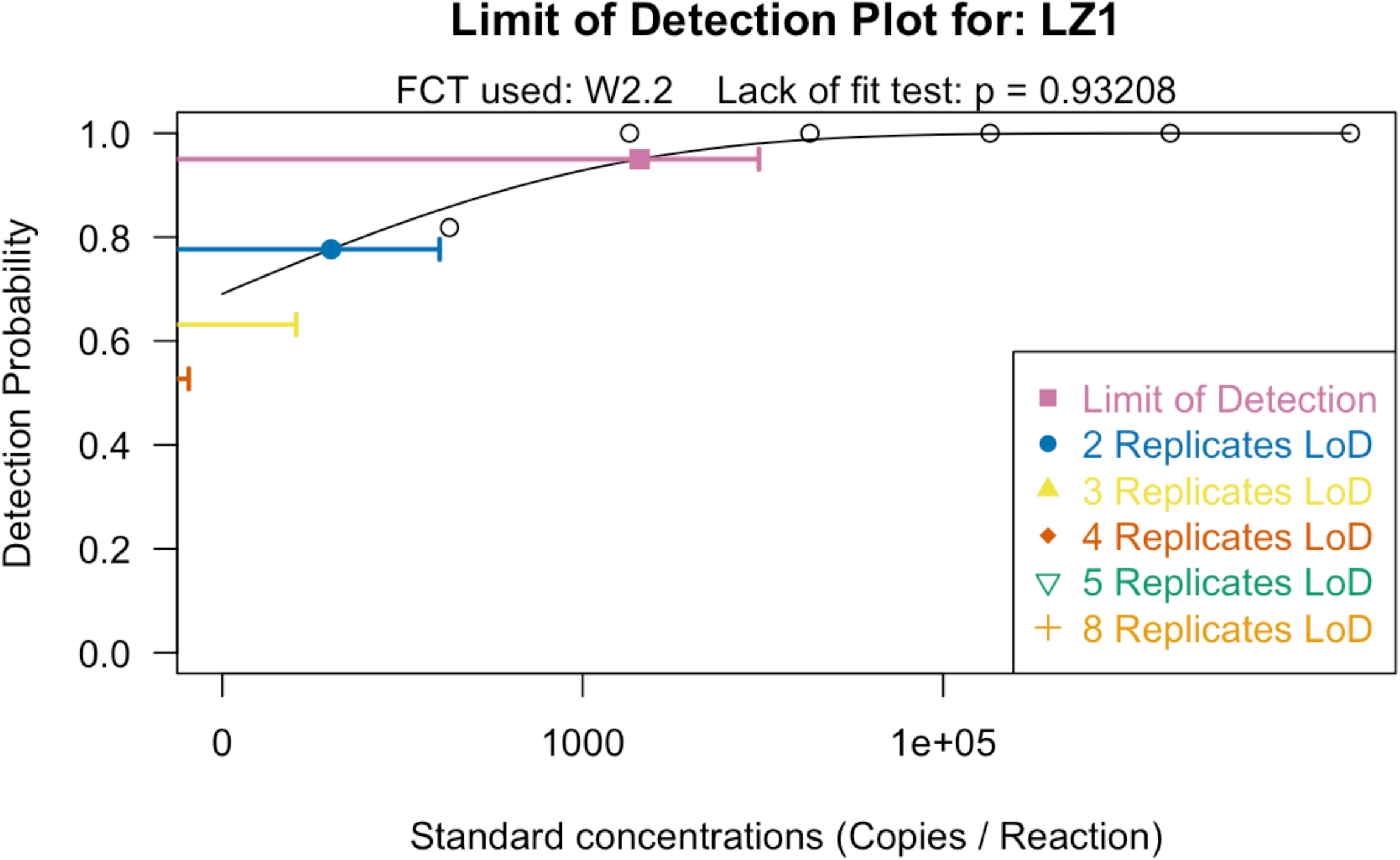
Limit of Detection (LOD) for LZ1 qPCR assay. X axis shows standard concentrations and y axis, the detection probabilities. Colours highlight the number of replicates required to achieve the LOD of 1820 copies uL^-1^. FCT is the logarithmic function used: Weibull Type II, 2 parameter function. This was used as it best fit the data. Method and R script from (60).

**Figure 8:**
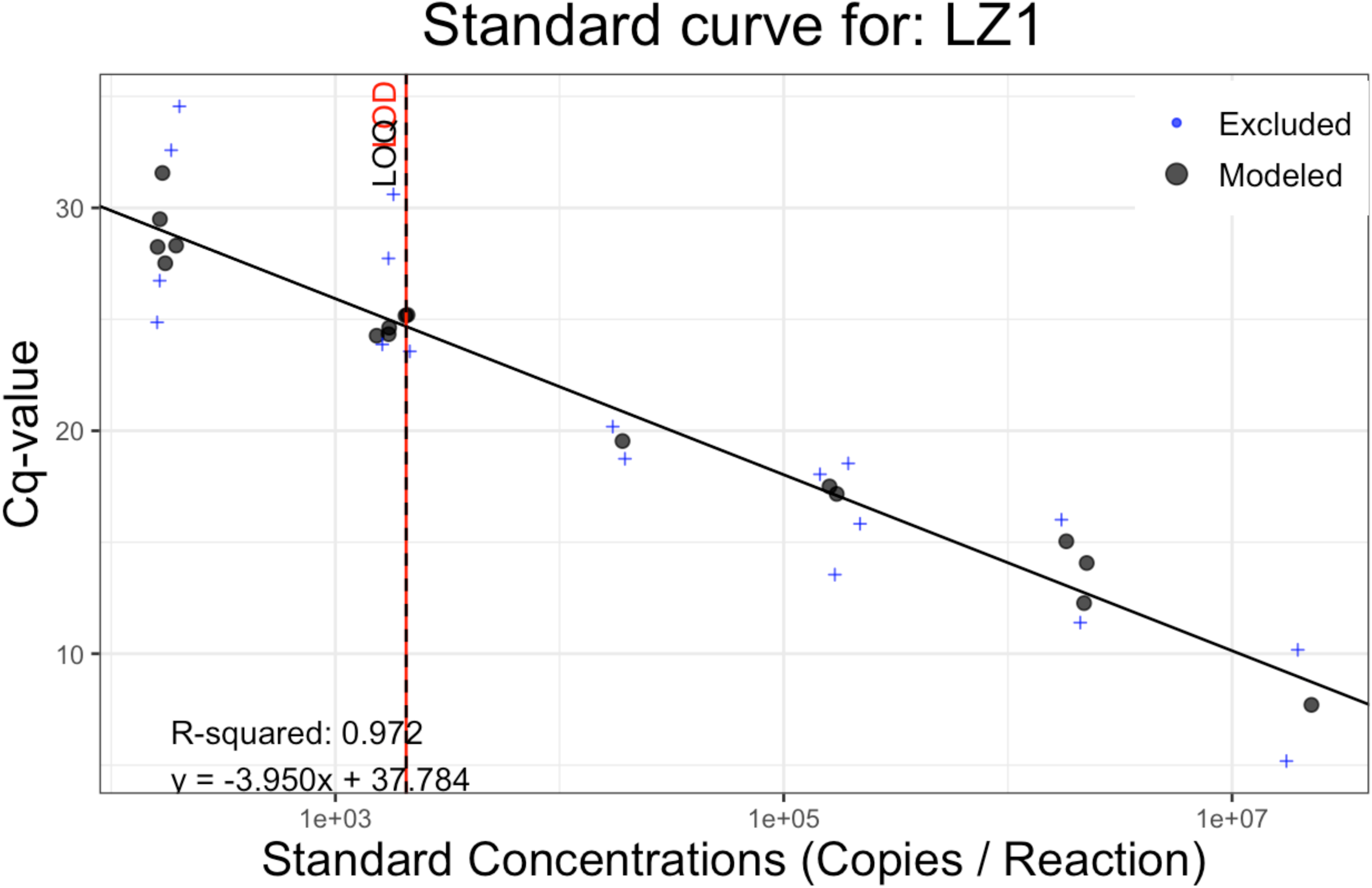
Standard curve for all runs of LZ1. Highlighting the limits of detection (black line) and quantification (red line).

Limits of detection of the LZ1 qPCR assay were tested by analysis of the detection probabilities of the standards at various concentrations using Weibull Type II, 2 parameter function to apply the best fit (60). The limits of detection of the analysis are 1820 copies uL^-1^. This correlates well with the results in Figure 5. The standard curve from all LZ1 analyses can be seen in Figure 10. This gives the Ct of a single copy (y-intercept) of 37.78. The limits of quantification can be improved by increasing the number of sample replicates, with an 80% probability of detecting 100 copies with >3 replicates.

LZ1 and LZ3 were then tested using SYBR green qPCR to further interrogate the performance of the primers on the test panel of organisms (Supplementary information 3). LZ2 was not tested further due to its poor end-point PCR performance. LZ1 gave Ct values around 21-22 for all the organisms positively identified using end point PCR. This equated to 1 x10^7^-10^8^ copies µL^-1^. All samples had the same concentration of DNA, 2 ng µL^-1^. Samples which tested negative in end point PCR had much lower Ct values - >32 and in some cases were undetermined relative to no template control (NTC). This was true for both primer sets. LZ3 had significantly lower Cts on organisms which tested positive using end point PCR. Typical Cts were 31-34, equating to a copy number between 1×10^4^-10^5^ copies µL^-1^. When comparing the results of the standard curves, LZ3 had slightly lower efficiency, 83.29% to 86.353% with a higher slope, 3.8 to 3.669. A perfect qPCR standard curve, with exponential amplification results in a slope of 3.3. However, LZ3 produced multiple peaks on a melt-curve analysis, indicating more than one product was generated and contributed to the fluorescent signal and was therefore ruled out as a suitable primer set for qPCR.

## Discussions

The need to develop improved microbial testing for water quality is evident from the lack of meaningful information that culture testing provides to water utilities. The rates of false positives and negatives have been well documented. The methods have been unchanged in over 100 years. Using molecular analyses like qPCR can improve the quantity and quality of the information that we have about the microbial environment in our drinking water systems, in addition to improving on the accuracy and speed of analysis. qPCR can be integrated with other molecular analyses, including testing for other pathogens, 16 rRNA sequencing and flow cytometry. This makes it an ideal method for investigation and risk assessment of water supplies.

### Phylogeny of Enterobacteriaceae

The *lacZ* database created for this study is the largest pool of sequences to-date gathered to assess PCR performance of coliforms. The phylogeny of the *lacZ* sequences in *Enterobacteriaceae* is complex, with *E. coli* species being particularly spread across the branches. This indicates a high degree of horizontal gene transfer for *lacZ*. However, analysis of 16S rRNA sequences also indicated that the amount of genetic distance was not significantly different between species in the database, and indeed much less so than the *lacZ* phylogeny. The concordant phylogeny demonstrates that the genus *Klebsiella* is most distant from other coliform and non-coliform *Enterobacteriaceae*, making it potentially more difficult to identify than other genera depending on the primer set. *Aeromonas* was shown to be more a closely related genus to coliforms than other non-coliforms with a *lacZ*. Overall, the phylogeny presented here demonstrates that the definition of coliforms as a group should be revised. They are often more closely genetically related to other *Enterobacteriaceae* which are not considered coliforms than they are to other species in the genus, due to the methodological conditions applied. Non-coliform species with a *lacZ* can also be associated with faecal contamination, like *Salmonella enterica* and *Vibrio cholerae*. There is a need to better define faecal indicator organisms in drinking water standards, which may focus less on the ability to metabolise ONPG at 37°C, and better understand risks to public health.

In silico results

The performance of primers found in previous research was mixed. The most employed primers in previous studies were 1675f/2025R (42–44, 46, 47, 59, 61, 62). These were the poorest performing primer of the sets tested. These primers were originally developed in 1990 and optimised in 1991. These results show that they target less than Primer set 2312f and 2548r and had >60% coverage across the database, scoring comparatively poorly compared to *de novo* primers. 2312f was reported to have an improved recovery than 1657f/2548r in a 2015 study (48). Primers 3153f and 2312f were also found to overlap but have different designations due to being generated using different databases and alignments. It should be noted that the position of all existing primers does not correspond to the positions noted within this expanded *lacZ* database. Overall, the *de novo* primers designed in this study performed significantly better than existing primers, potentially due to the introduction of redundant bases within PrimerProspector.

The most common genera recovered by all primers were *Escherichia, Enterobacter or Klebsiella* sequences. All primers also had perfect scores for *Shigella, Salmonella* or both. *Shigella* species are now defined within the *E. coli* taxon due their genotypic and phenotypic similarities (10, 58, 63). Other common non coliform genera recovered were *Aeromonas* sequences which were commonly returned by most primer sets, except 2312f, and 1551r. Primers 1411f and 3995r also did not return 0 scores for *Aeromonas*, but 3153f returned 3 sequences with 0. Primers 1441f, 3153f and 3995r scored more widely across all coliform genera, returning hits for: *Citrobacter; Cronobacter; Raoultella; Serratia* and *Pantoea*. No primer set scored 0 across all coliform genera. The recovery of more genera using these primer sets was offset by their reduced ability to recover the wide variation in *E. coli* and *Klebsiella* sequences across the database. 3153f and 3995r also returned *Vibrio* species with perfect matches, indicating that, despite their good score, they may not be discriminatory enough to identify only coliforms.

Analysis of variance was carried out on each distance matrix to test whether amplicons generated by coliform sequences were significantly different to non-coliforms (Group) and between amplicons from sequences from different species. No primer set generated amplicons with R^2^ > 20%, suggesting coliforms and non-coliform sequences do not have a significant difference in terms of distance (where R^2^ >0.2 p=<0.05). Speciation explained more variation with >99% variation in the amplicons in 1368f/1776r (FP1_RP1) explained by their species. This is potentially due to the primer’s proximity to an active site-Glu461, suggesting conservation of this motif has been driven by speciation, rather than horizontal gene transfer. This combination may then be a good candidate for recovery of coliforms with an active *lacZ* gene. FP3_RP3 & FP3_RP4 had >25% variation by Species and around 20% by Group. The best performing primer sets by Group were 2312f (FP5) in combination with 2548r, 3995r and 2546 (RP3) (p=0.01). As the amplicons generated by these had the most significant differences between coliforms and non-coliforms then this could suggest they provide the best discrimination of non-coliforms *in vitro* of the primers tested. Although this is not consistent with the other results. FP6_RP6 was the best performing *de novo primer* had almost none of the variation explained by either speciation or group.

### Top 3 primer sets in vitro

The primer results in vitro for the three sets tested were broadly supportive of the *in-silico* results. Existing primers, 2312f/2548r (LZ2) performed poorest in both the *in silico* and *in vitro* testing. This is despite these primers generating the highest amount of variance between non-coliform and coliform amplicons. This primer set failed to amplify several coliforms, including an environmental *E. coli* species, which is a key target organism for water utilities. These primers are formed from a combination of existing primers (forward: Molina et al, 2015, reverse: Bej et al 1990). Several studies have used the reverse primer to target coliforms in the environment (40–47). These primers were found to have low specificity and recover non-coliform organisms like *Salmonella* or *Shigella*. A study found that these primers also failed to target all known coliforms (9). This is consistent with the test panel results presented here, as these primers amplified the least number of coliforms. The forward primer from Molina et al was designed to improve upon the specificity of the original Bej et al primer, but here it does not prove to improve the ability to amplify representative organisms. For this reason, this primer set was not tested using qPCR.

LZ3, a more recently developed primer set had improved specificity than LZ2 (Zhang et al., 2015). These had better coverage than LZ2 but still amplified *Shigella* and *Salmonella* species (9, 48). Specificity testing of these primers was carried out using a small group of known coliforms in the original study, and they were developed using a small database of sequences (∼n=100). These had also not yet been tested in a qPCR assay until this work. 3153f/3995r (LZ3) performed best against the test panel as it did *in silico* and was the only primer to successfully amplify the target in all the coliform species. However, it did also amplify 2 non-coliform organisms as the *in-silico* results suggested it would. These primers were tested by the original designers against a wide test panel of organisms for identification purposes and these results support their use in identification of coliform bacteria. The large amplicon size generated by the primers makes it an ideal set for sequencing of amplicons.

*De novo* primers, LZ1 performed better than LZ2. This had a smaller amplicon size and was able to identify all coliforms but a single laboratory *Enterobacter* species. Although it did amplify an environmental *Enterobacter*, suggesting its ability to recover organisms in this genus. It also recovered the species *Aeromonas veronii*, a non-coliform which was picked up by all primer sets. This is interesting as when initially cultured, the colony appeared yellow, which meant it was counted as a presumptive coliform based on the MLGA testing. This species was also closely related to coliform species on the *lacZ* phylogeny, indicating it may always be difficult to distinguish from coliforms. It should be noted that identification of species was based on sanger sequencing of partial 16S rRNA amplicons, sequencing of a longer length may be required to verify the classification. However, this example and the significant amount of variation not explained in the PHYANOVA results, demonstrates that classification of coliforms is difficult, no matter what the methodology. The group may need to be widened or re-considered to include other species which may also be indicative of faecal contamination. LZ1 out-performed LZ3 in the qPCR assay on the test panel organisms. As the DNA concentrations of the test panel were standardised to 2 ng uL^-1^, the primer sets should have resulted in equivalent Ct values, however LZ3 had Ct values of around 32-34 compared to 21-22 of LZ1. The efficiency (86.53%) and slope (3.669) of LZ1 were within acceptable limits for an effective qPCR assay, but LZ3 had lower efficiency. LZ3 also produced multiple peaks on a melt curve analysis. LZ1 were therefore the best primer set for testing *in vivo*.

Sequencing of *lacZ* from more environmental organisms would allow further optimisation and specification of primer sets to improve recovery across the coliform group. In addition, developing a database of *lacZ* sequences can help water authorities understand commonalities in coliforms detected in the environment, helping to narrow down their source and limit impacts to public health. *Aeromonas, Salmonella* and *Shigella* species are common false positives in coliform testing, meaning confirmatory procedures must be exhaustive, e.g.: oxidase testing; indole production and growth at 44°C (8, 64, 65). Combining culture testing with a PCR confirmatory step can help reduce false positives and negatives, as well as speed up analysis time by up to 24 hours. *Salmonella* and *Shigella* are also a significant water-borne health risk and being able to track these via sequencing. Shigellosis is considered one of the most significantly underreported diseases associated with drinking water in the European Union alongside Giardiasis (66).

Based on these results, no primer set is likely to be able to discriminate all non-coliform *lacZ* sequences. This is concurrent with previous studies proposing a high level of horizontal gene transfer in the evolution of *lacZ* as well as the phylogeny presented here (Bej et al., 1991b, 1990b; Fricker and Eldred, 2009; Fricker and Fricker, 1994; Horáková et al., 2006; Molina and Lowe, 2012; Zhang et al., 2015). Other studies have highlighted the same problem with other targets like *tufA; gap; ompA; rpoB* and *infB* (10, 62). This emphasises false positives/negatives are a problem when analysing coliforms both *in vitro* or *in vivo*, and the group’s lack of phenotypic and genotypic similarities make these only a group in terms of their ability to be cultured using specific growth media, often under stress. This is furthermore emphasised by the lack of an available qPCR probe site. Potential probe sites were assessed using PrimerProspector but no site in addition to the primer set was specific enough to develop a TaqMan® probe. This meant that the qPCR assay had to employ SYBR green rather than TaqMan® for fluorescence. TaqMan probes are more specific than SYBR green due to their specific binding. However, as the LZ1 amplicon is quite small (150bp) the interference from partial binding of SYBR green should not be as significant as it may be in larger amplicons.

Sensitivity testing and method comparison.

When tested against other typical water quality tests like MLGA membrane filtration and flow cytometry, LZ1 qPCR performed less well than culture testing at low concentrations (<1 x 10^3^ 100 mL^-1^. The experimental detection limit from this study was equivalent to 200 cells per reaction and a limit of detection of 1820 cells µl^-1^. However, it should be noted, that in this study while 100 mL of water was filtered, equivalent of only to 20 mL of water was added to the qPCR reaction, as not all the DNA is used. As such our qPCR method detected 200 cells per 20 ml or 10 cells per 1 mL. Therefore, by increasing sample volume and the number or replicates, the recovery and accuracy at low concentrations can be significantly improved. Despite this, qPCR results generally correlated well with the MLGA results (R^2^=0.88) in the pure water samples. The drinking water samples had less correlation, indicating that background DNA may amplify the signals, particularly in the lower concentration samples, or that there were background copies of coliforms in the water used. The blank sample (no *E. coli* added) had 2.62 log copies in the original sample. However, this is lower than the LOD and LOQ. To improve the LOQ to around 2 logs, an increased number of replicates could identify whether this is low-level coliform DNA present in the drinking water, or background signal. MLGA result was 0 cfu 100 mL for this sample. The higher concentrations (>1 x 10^3^ 100 mL^-1)^ were much closer in terms of accuracy, indicating that filtration of a higher volume will also improve quantification. This is also supported by a recent study, where filtration of large volumes improved recovery of opportunistic pathogens in drinking water systems from 77-83%. Many of these were found to be viable despite the presence of free chlorine (67). This further emphasises the need for improved coliform detection.

One important consideration which must be addressed, is the viability of the *E. coli* cells. When enumerating the cells to carry out the serial dilution in this experiment, a total cell count was made using Flow Cytometry. This count includes both alive and dead cells. Similarly, the qPCR assay cannot discriminate between a living cell and extra cellular DNA. This may explain the level of background signal in the drinking water samples which was not evident in the MLGA. This may also explain why the flow cytometry counts were much more similar between drinking water and pure water samples compared to the culture testing.

Understanding the viability of cells is crucial for water utilities, particularly when assessing the risk to public health. Therefore, a pre-treatment with propidium monoazide is proposed when using this assay to reduce interference from partial or dead cells (68).

At low *E. coli* cell abundance and sample volumes, culture testing is both more accurate and sensitive at returning low level viable organisms. This is in part due to all the sample being used for the analysis. Compared to flow cytometry, where the small analysis volume (50 µL) makes sensitivity and accuracy poor at concentrations <1 x 10^2^ 100 mL^-1^. The advantages of a qPCR assay are that significantly larger volumes of water can be analysed via filtration. Further, this work has shown that by increasing sample replicates the limits can be significantly reduced increasing the probability of quantifying a low abundant template. DNA extraction and qPCR can be run in a single day, whereas culture testing takes at least 24 hours, for a negative result and at least a further 24 hours to run confirmatory testing on any positives. A further advantage of filtration of a large volume sample, is that it can be used to test against a variety of qPCR targets. So, a sample can be tested for coliforms, *E. coli, Salmonella* and other indicator and pathogenic organisms to risk assess a particular water supply. This DNA can also be used to carry out metagenomics or 16S rRNA amplicon sequencing, to provide a more overall picture of the microbiome.

### Application of *lacZ* qPCR for drinking water

Based on these results, a qPCR assay is unlikely to replace routine microbial culture testing for coliform bacteria. That does not mean that the assay doesn’t serve an important function to water utilities. Water system risk assessments are an important component of managing water quality and are recommended by the World Health Organisation (WHO). Their guidance states that the absence of coliform bacteria is not sufficient to demonstrate the safety of a drinking water supply, a robust risk assessment approach is also required (69). QMRA is such an approach (70, 71). LZ1 qPCR could be a critical component of a Quantitative Microbial Risk Assessment (QMRA) approach to managing drinking water quality. This is a multi-step risk assessment process which first seeks to quantify the risk of each possible pathogenic microbe in the source, and the risk post treatment. The likelihood and volume of consumption for each pathogen is then calculated per individual. This helps utilities target investment and improvement to supplies where the risk of pathogens is higher. Countries like Australia and the Netherlands have already adopted a QMRA approach to their drinking water supplies with great success (70).

The large volume that can be analysed via qPCR is much more representative than the typical 100 mL sample taken for routine coliform testing. This is an infinitesimal fraction of the volume produced by a typical treatment plant. Furthermore, the sample is taken at a point in time, by filtering a larger volume, sampling can be undertaken over a longer period, giving a better representation of a plant’s performance. This is like typical cryptosporidium analysis, where large volumes are filtered over a period of several hours to days (UK Government, 2010). Also, like cryptosporidium testing this analysis could be undertaken during periods of high-risk, when treatment processes are sub-optimal or raw water conditions have deteriorated. This would allow water utilities to demonstrate to their regulators and customers the impact of these issues on the wholesomeness of the supply.

Another significant benefit of qPCR testing for coliforms is to further analyse their impacts to public health. As faecal indicators, total coliforms are known to be poor and current approaches to QMRA are limited to estimating pathogens in the source based on literature, and therefore may significantly over or underestimate concentrations. Combining LZ1 qPCR alongside assays for *E. coli* specifically and other enteric pathogens like *Salmonella* and Campylobacter can further explore the health risks inherent in a drinking water supply. Non-enteric pathogens like *Legionella* and *Mycobacterium* are also of emerging risk to water utilities. Culture and speciation of *Legionella* is time consuming, and no available methodology is available to culture *Mycobacterium*, but qPCR tests exist to quantify both. Therefore, investing in methods for filtration of large volumes of water, DNA extraction, qPCR analysis can provide significant benefit to a water utility. These benefits are multiplied when qPCR analysis is combined with other molecular analyses like flow cytometry and 16S rRNA amplicon sequencing. These in combination can provide significantly more information than culture testing alone can provide.

## Conclusions

Coliform bacteria remain the primary indicator of microbial contamination of drinking water supplies for many water regulators, including Europe, the U.K., and the U.S.A. This paper further challenges the reliance that regulators place on these as a group of bacteria. These are broadly phylogenetically unrelated and don’t form linear speciated groups as culture testing would imply. Moreover, they provide very little information on the overall quality of the water and are difficult to isolate the cause. Using qPCR to test for coliforms as well as other pathogens in combination with more general molecular techniques like flow cytometry and 16S rRNA sequencing, will allow utilities to test microbial removal more thoroughly and isolate contamination more quickly than traditional snap sampling. There are a few existing primers and assays designed to target coliforms in drinking water, however the performance of these has been mixed. Using a larger database of sequences to refine target sequences has improved the recovery of coliform organisms using PCR, however false positives and negatives remain problematic, due to the high amount of phylogenetic and phenotypic variation among the group. This further highlights the issues in relying on coliforms alone as an indicator of public health. LZ1 is the best option for enumerating coliforms via qPCR due to its small size and relative efficiency compared to the other primers tested.

## Acknowledgements

Funding: CT. Scottish Water Industry Funded PhD studentship; by EPSRC award EP/V030515/1, and EP/W037270/1 (UoS) and EP/W037475/1 (UoG), and a Royal Academy of Engineering-Scottish Water Research Chair (RCSRF171864) awarded to CJS.

